# High-Pressure NMR Reveals Volume and Compressibility Differences Between Two Stably Folded Protein Conformations

**DOI:** 10.1101/2020.07.29.227322

**Authors:** Xingjian Xu, Donald Gagné, James M. Aramini, Kevin H. Gardner

## Abstract

Proteins often interconvert between different conformations in ways critical to their function. While manipulating such equilibria for biophysical study is often challenging, the application of pressure is a potential route to achieve such control by favoring the population of lower volume states. Here, we use this feature to study the interconversion of ARNT PAS-B Y456T, which undergoes a dramatic beta-strand slip as it switches between two stably-folded conformations. Coupling high pressure and biomolecular NMR, we obtained the first quantitative data testing two key hypotheses of this process: the slipped conformation is both smaller and less compressible than the wildtype equivalent, and the interconversion proceeds through a chiefly-unfolded intermediate state. Our work exemplifies how these approaches, which can be generally applied to protein conformational switches, can provide unique information that is not easily accessible through other techniques.

## INTRODUCTION

The energy landscape theory of folding states that proteins sample many conformations before reaching the native state ^[1, 2]^. Although the native conformation is usually the lowest free energy state under physiological conditions, proteins can be trapped in stably-folded local minima that are markedly different from the native conformation ^[3-5]^. Transitions between conformations, which can be spontaneous or triggered by changes in environmental conditions (e.g. pH, temperature, light) or ligand binding, are often critical for shifting proteins between functionally inactive and active forms ^[3-7]^. Such shifts can range from simple local rearrangements to ‘metamorphic’ proteins, which reversibly adopt different stable folds in different environmental conditions ^[8-10]^. However, identifying and characterizing alternative conformational states of proteins can be challenging, as they are often high-energy states and sparsely populated ^[11, 12]^. Several techniques currently allow quantitative characterization of conformational sub-states despite challenges introduced by low population, particularly solution nuclear magnetic resonance (NMR) techniques such as relaxation dispersion experiments ^[11-14]^.

To aid in characterizing such excited states, it is routine for one to manipulate their populations by adding small molecule ligands or changing experimental conditions, such as pH or temperature ^[11, 15-17]^. Increasingly, pressure has also been utilized to control conformational equilibria, aided by the introduction of commercially-built pumps and sample cells compatible with NMR and other biophysical instrumentation ^[18-20]^. Pressure directly affects such equilibria by acting on differences in the partial volumes and compressibilities of different conformers, with increasing pressure favoring those with lower volumes ^[21]^. Given that unfolded proteins are generally smaller than their folded forms, pressure studies have been particularly useful at investigating protein unfolding reactions, which typically occur above 2000 bar under native conditions ^[20-25]^. However, the application of sub-unfolding pressures can cause proteins to shift populations among multiple folded states ^[18, 26-28]^, providing an easy way to manipulate protein conformational equilibria to facilitate their study.

These advantages of pressure-NMR led us to examine its applicability to characterize a protein domain which slowly exchanges between two folded states via an unusual β-strand slip ^[29, 30]^. This system is derived from the human ARNT (Aryl hydrocarbon Receptor Nuclear Translocator) protein, a bHLH-PAS (basic helix-loop-helix / Per-ARNT-Sim) transcription factor which binds a variety of partners via its two PAS domains, PAS-A and PAS-B ^[31-35]^. PAS domains adopt a mixed α/β fold, with helical and sheet layers often encapsulating internal cavities between them ^[31, 32, 34]^. In many cases, these internal cavities provide binding sites for natural or artificial regulatory molecules ^[36]^. For ARNT PAS-B, 105 Å^3^ of internal cavities are seen in the crystal structure, suggesting that small molecules may bind there and control the role of ARNT in several signaling pathways ^[29, 32, 37, 38]^.

We have previously reported that the ARNT PAS-B β-sheet can surprisingly adopt an alternatively-folded conformation in certain settings. For example, a Y456T point mutation at a site preceding the final β-strand enables the domain to adopt a new conformation that coexists in a 1:1 equilibrium with the WT fold ^[29, 30]^ (Fig. 1). Additional mutations to nearby residues (F444Q/F446A/Y456T, hereafter called the TRIP variant) stabilized this alternative conformation, letting us characterize the new structure and show that it differs from the WT conformation by a three residue slip of central Iβ-strand of the domain’s five-stranded β-sheet. This slip inverts the topology of the Iβ-strand and isomerizes the neighboring N448-P449 peptide bond, collectively abolishing the domain’s ability to interact with some other binding partners ^[29]^. Notably, the threonine side chain introduced by the Y456T mutation fills an internal cavity ^[29, 30]^, leading us to suspect that interconversion process could be manipulated by pressure perturbations ^[18, 39-42]^ to obtain thermodynamic and kinetic information complementary to our prior structural studies.

**Figure 1.**
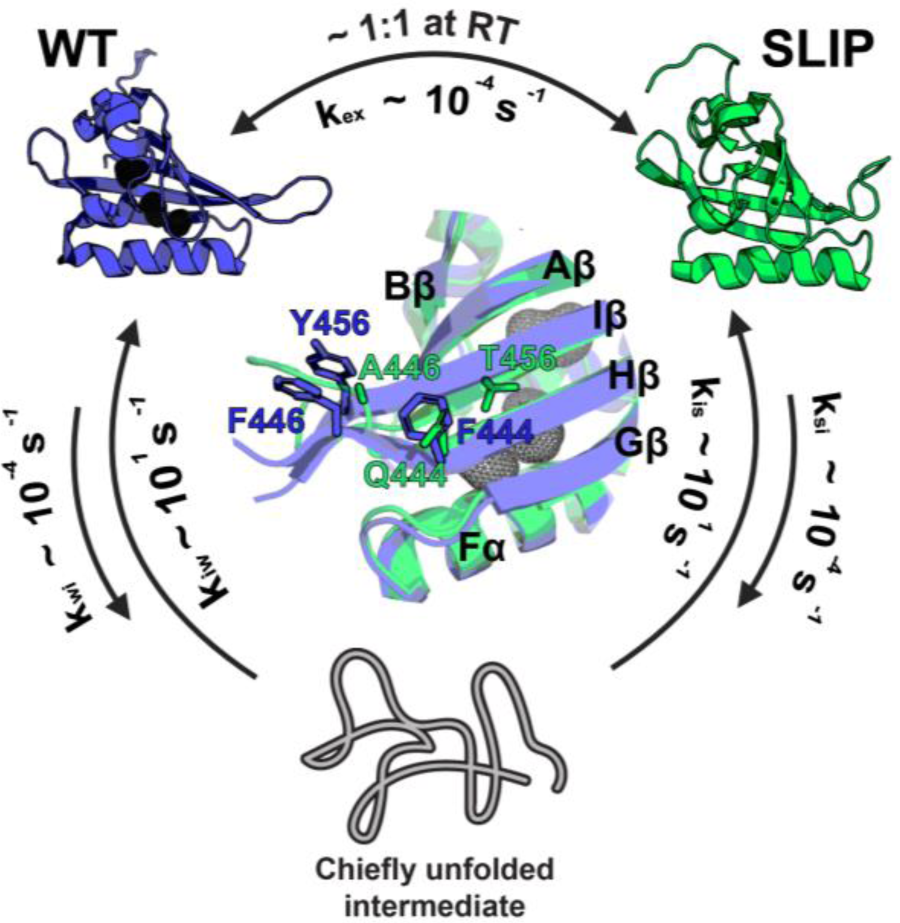
Current model of ARNT PAS-B Y456T interconversion between the WT and SLIP conformation. Prior work demonstrated that this protein interconverts between the folded WT and SLIP conformations on the same timescale as the unfolding rates of either conformer, implying a slow transition process requiring a chiefly unfolded intermediate state. The two conformations are in 1:1 equilibrium at room temperature, but lower temperature favors the SLIP conformation. The F444Q/F446A/Y456T variant (TRIP) locks the protein in the SLIP conformation, structural comparison between WT and TRIP (WT slate blue, SLIP lime green) is displayed in the middle, with mutation sites labeled.

Here we report results from high-pressure solution NMR studies of ARNT PAS-B Y456T, letting us for the first time obtain quantitative measurements of several thermodynamic and kinetic parameters of the WT:SLIP interconversion. We found that the WT conformation is not only larger in volume than SLIP, but also more compressible as perhaps due to the cavities unique to the WT conformation ^[29, 32]^. Additionally, we tested a hypothesis that WT:SLIP interconversion proceeds through a chiefly unfolded intermediate state, previously suggested from the similarities of interconversion and unfolding rates ^[30]^. We found that pressure increases the rate of interconversion, letting us measure both the activation volume and compressibility ^[30]^, both validating this model and allowing us to quantitate how unfolded the transition state is. Finally, we demonstrated that both residue-specific compressibility and pressure-dependent chemical shift changes can predict whether residues are near cavities, providing insights into locations of such cavities within the protein. Taken together, our approach exemplifies the ability of high-pressure NMR to enable thermodynamic and kinetic analyses of protein conformational transitions that are otherwise inaccessible.

## RESULTS AND DISCUSSION

### Equilibrium Analyses: Pressure Dependence of ARNT PAS-B Conformation Equilibrium

We previously determined the solution and crystal structures of WT ARNT PAS-B and the solution structure of the F444Q/F446A/Y456T (TRIP) variant ^[29, 31, 32]^. By comparing the average void volumes of the two solution structure ensembles with ProteinVolume ^[43]^, we found that the SLIP conformation had approximately 600 Å^3^ smaller void volume than WT, leading us to anticipate that the equilibrium between these conformations might be pressure sensitive. To test this possibility, we recorded ^15^N/^1^H and ^13^C/^1^H HSQC spectra of ARNT PAS-B Y456T at increasing pressures (20, 250, 500, … up to 2500 bar). Overlaying these spectra (Fig. 2A, S1), we noticed that signals from all three (^1^H, ^15^N, ^13^C) nuclei showed pressure-dependent chemical shift and intensity differences between the two conformations, as predicted from the volume differences between them. In general, we observed that increased pressure decreased the intensities of WT signals while concomitantly increasing SLIP signal intensities with good reversibility (Fig. S2), confirming the smaller calculated volume of the SLIP conformation ^[30]^.

**Figure 2.**
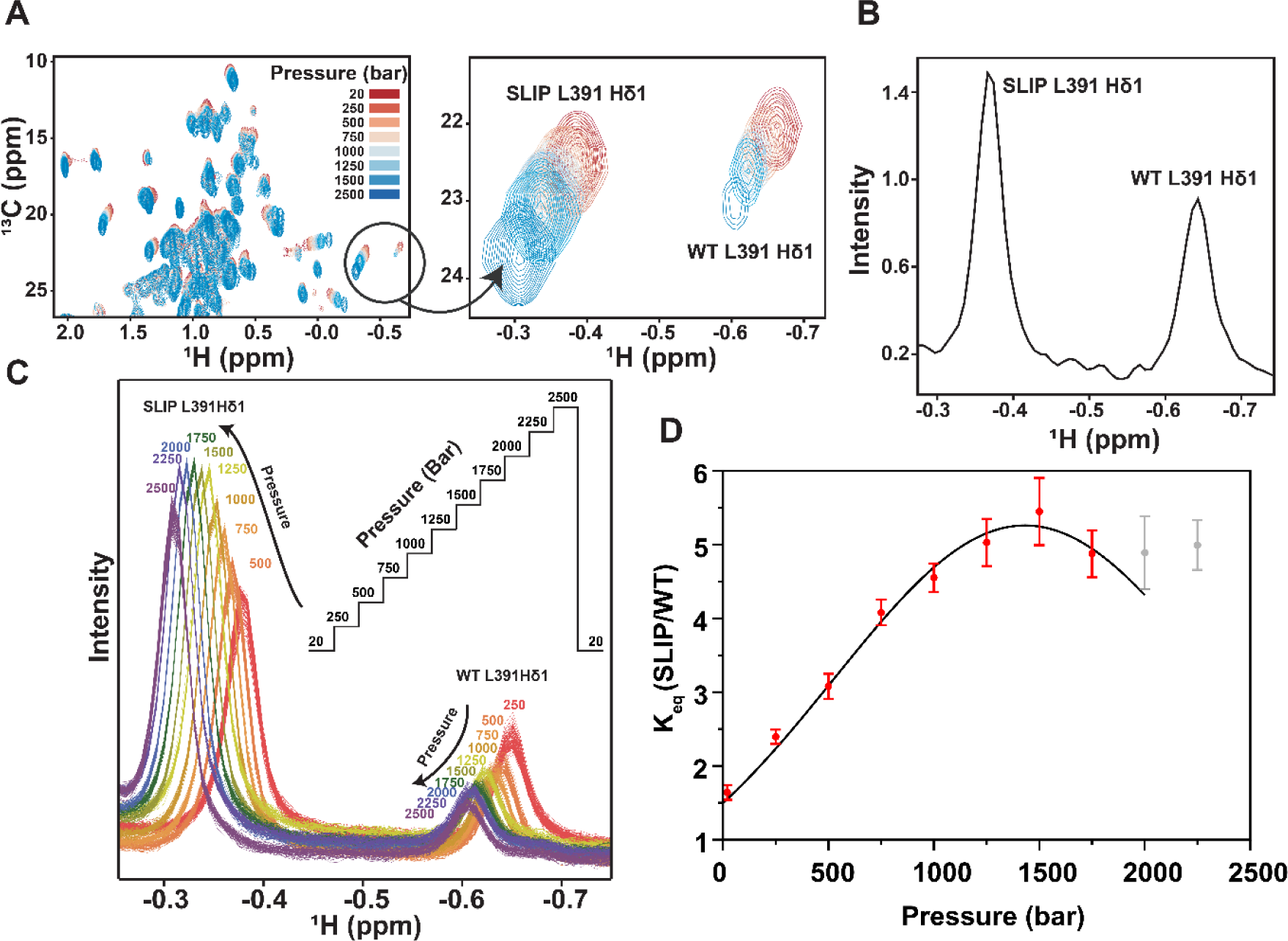
NMR evidence for pressure-induced equilibrium shifts between WT and SLIP conformation of ARNT PAS-B Y456T. A) Methyl region of ^13^C/^1^H HSQC spectra of ARNT PAS-B Y456T recorded after equilibration between 20-2500 bar. Pressure-dependent intensity changes at several sites, including the L391 δ1 methyl group, reflect a shift in the SLIP:WT equilibrium. B) ^13^C-edited 1D ^1^H spectra of L391 Hδ1 equilibrated at 278.1K and 20 bar. *K*_*eq*_ (SLIP: WT) is approximately 1.5, as assessed by the areas of the two conformer-specific peaks. C) ^13^C-edited 1D ^1^H NMR of L391 Hδ1 recorded at pressures between 20-2500 bar as shown in inset. Spectra were collected during sample equilibration at each pressure. D) Fitting pressure dependence of equilibrated *K*_*eq*_ from datasets shown in C. *K*_*eq*_ above 1750 bar are not included for the fitting as another SLIP peak moves close to SLIP L391 δ1, interfering with the baseline and leading to overestimation of the SLIP population.

To characterize the thermodynamics and kinetics of the WT:SLIP conformational interconversion, we acquired a series of ^13^C-edited 1D ^1^H NMR spectra as the system equilibrated after pressure changes. We have previously shown that the upfield-shifted L391 Hd1 methyl signal of both WT and SLIP conformations are well resolved in multiple types of NMR spectra, likely due to its location adjacent to a cavity only found in the WT conformation (Fig. S3) ^[30]^. Using this methyl group as a probe, we determined the equilibrium constant *K*_*eq*_ (=[SLIP]/[WT]) at different pressures (20 to 2500 bar) by measuring the ratios of the peak integrals corresponding to these two conformations (Fig. 2B, C).

From these data, we extracted the pressure dependence of *K*_*eq*_, noting that it monotonically increased until approximately 1500 bar. Beyond 1500 bar, we observed that it remained approximately constant, showing that *K*_*eq*_ does not scale exponentially with pressure and correspondingly that the free energy Δ*G* is not linearly dependent on pressure p (Eq. S1 and Fig. 2D), necessitating the inclusion of a nonlinear compressibility term for our analyses. Fitting these data to obtain differences in free energy between the two conformations at ambient pressure (referenced to 1 bar, Δ*G*^0^), volume (Δ*V*^0^), and compressibility (Δ*βV*^0^) (Eq. S3), we found that the SLIP conformation was 40.5 ml/mol smaller than the WT conformation, reasonably agreeing with the ca. 105 Å3 (= 63 ml/mol) internal cavities unique to the WT structure. Additional support for the pressure-driven loss of these cavities came from the measurement that the WT conformation was more compressible than the SLIP by 0.0285 ml/(mol bar). We attribute both the volume and compressibility differences to the collapse of internal cavities and increased packing accompanying the WT to SLIP transition [29, 32].

### Pressure Jump Kinetic Analyses: WT and SLIP Interconvert by Transitioning into an Intermediate State with Small Volume and Compressibility

To complement the above equilibrium pressure analyses, we used a pressure jump protocol to obtain complementary kinetics measurements. To do so, we jumped the sample pressure from 20 bar to various higher values, acquiring 1D ^13^C-edited ^1^H NMR spectra as the system re-equilibrated and measuring the populations of the two conformations by the area of the respective L391 Hδ1 signals (**Fig. 3A**). Notably, we conducted these studies with commercially-available instrumentation capable of completing even the largest of these jumps within 30-60 sec, making it possible for us to perform the kinetic analyses due to the slow kinetics of the interconversion process (**Fig. 3A**). Custom-built, NMR-compatible pressure systems have recently been reported which are capable of such jumps approximately 100-fold faster ^[23, 24]^.

**Figure 3.**
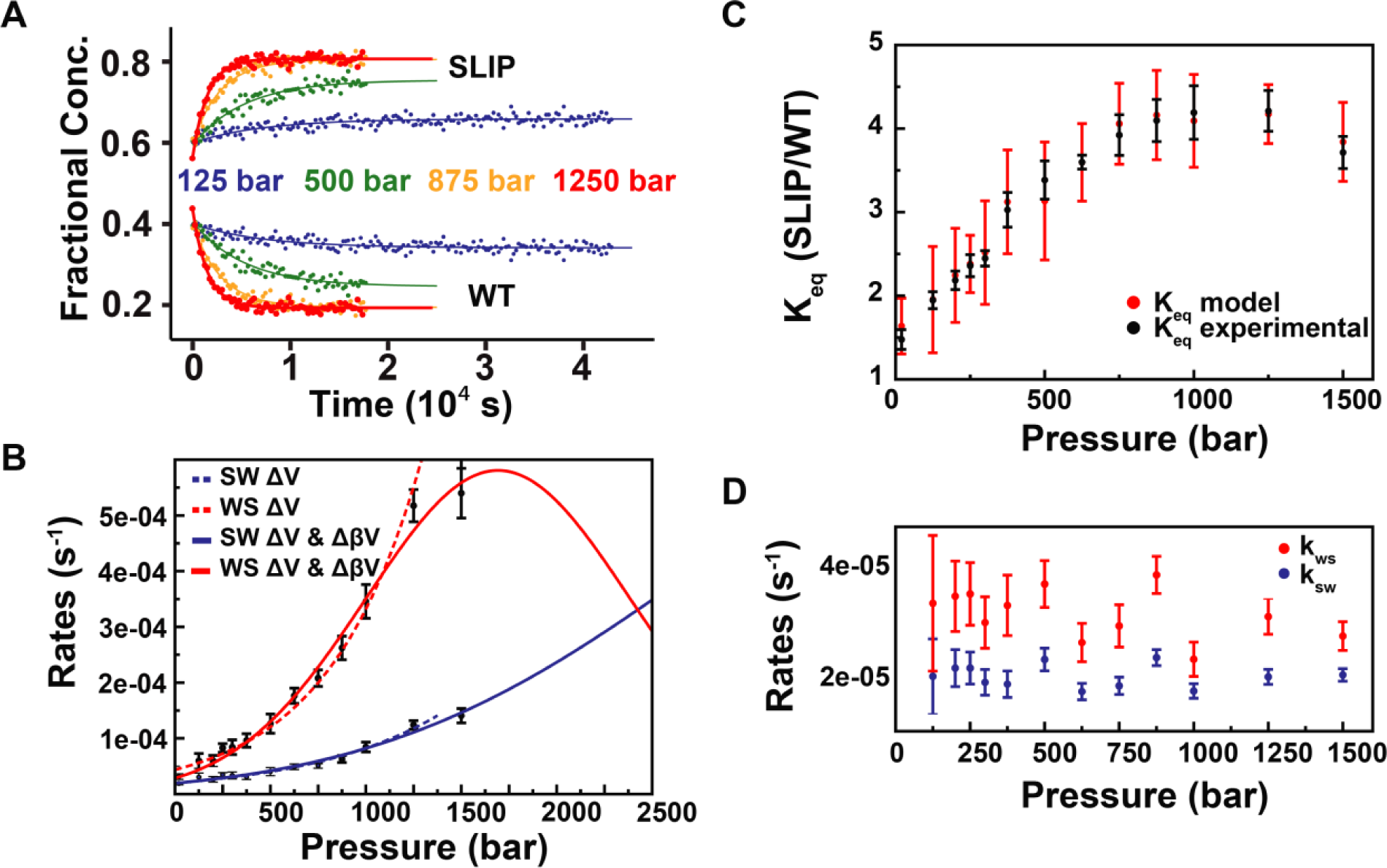
Kinetic analysis of the ARNT PAS-B interconversion. A) Examples of independent fits of the rate constants using the L391 Hδ1 ^1^H methyl signals at the indicated pressures. Total population of the WT and SLIP was normalized to 1 for each analysis. B) Fitting of the decoupled rates (*k*_*sw*_ in blue, *k*_*ws*_ in red) as functions of pressure. Two models (without and with compressibility implemented) were used to fit the experimental data, using dashed (solid) lines to indicate fits without (with) compressibility included. C) Comparison of fit model (red) to experimental data (black). Equilibrium constants from the model are calculated from the rates derived from the kinetic analysis. D) Decoupled rates of relaxation. Each rate pair is plotted against the pressure the system was relaxed from. Relaxation rates appears to be independent of the initial pressure.

As an initial control of this approach, we extracted the *K*_*eq*_ values from these direct jump experiments and fit them for the same parameters mentioned in the equilibrium analyses (**Fig. S4**). This yielded a Δ*V*^0^ of -46.1 ml/mol, and a Δ*βV*^0^ of -0.0433 ml/(mol bar) (**Table 1**), comparable to values we noted above. While direct pressure jumps may induce convergence of *K*_*eq*_ at a slightly lower pressure, as indicated by the bigger difference in compressibilities between the two states, we otherwise believe that these values are similar despite different experimental routes.

**Table 1.** Summary of the kinetic and thermodynamic parameters derived from pressure-induced interconversion and unfolding of ARNT PAS-B variants

| Thermodynamic Analyses |  | Kinetic Analyses |  |
| --- | --- | --- | --- |
| SLIP – WT, step jumps<sup>a</sup> | | with $\Delta\beta V$ | |
| $\Delta V^0$ (ml/mol) | $-40.5 \pm 0.47$ | $\Delta V_{SW}^\ddagger$ (ml/mol) | $-37.6 \pm 3.0$ |
| $\Delta\beta V^0$ (ml/(mol bar)) | $-0.0285 \pm 0.00065$ | $\Delta V_{WS}^\ddagger$ (ml/mol) | $-82.4 \pm 7.4$ |
| SLIP – WT, direct jumps<sup>b</sup> | | $\Delta\beta V_{SW}^\ddagger$ (ml/(mol bar)) | $-0.00875 \pm 0.0034$ |
| $\Delta V^0$ (ml/mol) | $-46.1 \pm 0.41$ | $\Delta\beta V_{WS}^\ddagger$ (ml/(mol bar)) | $-0.0487 \pm 0.0066$ |
| $\Delta\beta V^0$ (ml/(mol bar)) | $-0.0433 \pm 0.00067$ | without $\Delta\beta V$ | |
| Unfolding (U - F) | | $\Delta V_{SW}^\ddagger$ (ml/mol) | $-32.8 \pm 0.62$ |
| WT (ml/mol) | $-126.3 \pm 3.1$ | $\Delta V_{WS}^\ddagger$ (ml/mol) | $-59.0 \pm 1.38$ |
| TRIP (ml/mol) | $-83.3 \pm 2.3$ | | |

Uncertainties were estimated by bootstrapping. Random noises with mean of 0 and variance of the standard error were added to the raw data and fit repeatedly for $n \geq 30$ times to determine the 95% confidence interval. <sup>a</sup> step jumps – jumping pressure incrementally with 250 bar steps. <sup>b</sup> direct jumps – jumping pressure directly from 20 bar to various high pressures.

To analyze the kinetics of the interconversion process, we assumed that a two-state model could be applied by our prior observation that the intermediate state of ARNT PAS-B Y456T is only transiently visited during interconversion ^[30]^. We validated this assumption by establishing that the sum population of the two states remained constant at pressures below 2500 bar, suggesting the intermediate is not accumulating (**Fig. S5**). In the two-state transition model, the apparent exchange rate is the sum of the two rate constants corresponding to the transition from SLIP to WT and vice-versa (*k*_*sw*_, *k*_*ws*_) ^[44]^. To obtain the pressure dependence on both rates, we initially fit the apparent exchange rate vs. pressure to a biexponential equation (**Eq. S4**), to yield the initial rates at 1 bar (*k*_*sw*0_ and *k*_*ws*0_), the activation volumes (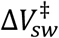 and 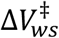), and the activation compressibilities (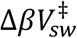 and 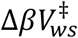). To avoid needing to simultaneously fit six parameters, we decoupled the two rates by solving them numerically under different pressures as described in Materials and Methods (**Eq. S5, S6**; **Fig. S6, S7**). This allowed us to independently fit each rate as a function of pressure with three parameters (**Eq. S7, S8**; **Fig. 3A, B**), summarized in **Table 1**.

From these analyses, we observed negative activation volumes and compressibilities for both transitions, implying that they proceed through an intermediate state which is both smaller and less compressible than the starting conformations. Moreover, the magnitude of the volume barrier is bigger for the WT to SLIP transition, with a 44.8 ml/mol difference in activation volumes, consistent with the 40.5 ml/mol volume difference obtained from the thermodynamic analysis. Taking the ratios of the two rate constants (*k*_*ws*_/*k*_*sw*_) at different pressures yielded equilibrium constants matching experimental data (**Fig. 3C**). Since both activation volumes were negative, the transition rates were correspondingly accelerated under pressure. Interestingly, the activation volume for the WT to SLIP transition (−82.4 ml/mol, or 137 Å^3^/molecule) is strikingly similar to the cavity size of the WT conformation. The SLIP to WT transition, in contrast, required a much smaller activation volume (**Table 1**). While the pressure-induced volume change of a system is a combined effect from compressing both protein and solvent ^[20]^, we found a positive correlation between cavity size and activation volume. From these data, we postulate the transition between the two confirmations requires the protein to collapse its internal cavities and voids to reduce its total volume, consistent with substantial unfolding that we quantitate below. This claim is further supported by Roche *et al*.’s ^[45]^ recent work highlighting that pressure-driven protein unfolding is a result of cavity elimination.

Additional support for an unfolded transition state is provided by the negative activation compressibilities, as the replacement of interatomic contacts with hydration shells during unfolding will substantially reduce compressibility ^[46]^. Notably, as compressibility is small and neglectable at lower pressures ^[18]^, we were able to fit the sub-1000 bar data points to a model without compressibility, leading to the same observation that the SLIP conformation is smaller than the WT conformation, albeit to a lesser extent (**Fig. 3B**; **Table 1**).

### Pressure Induced Interconversion is Reversible and Provides Thermodynamic Information on Transition State

To validate the reversibility of the pressure-jump experiments, we recorded ^13^C-edited 1D ^1^H NMR spectra as samples were dropped from higher pressures down to 20 bar. From these data, we extracted the two rate constants *k*_*ws*_ and *k*_*sw*_ and plotted them against the pressures prior to the relaxation (**Fig. 3D**). For all of the pressures we used for extrapolating the rates (from 125 to 1500 bar), the relaxation rates remained the same and the corresponding [SLIP]/[WT] equilibrium constants were the same as we measured in equilibrium studies at 20 bar. These results, together with our comparison of ^15^N/^1^H HSQC spectra recorded before and after pressure-jump experiments (**Fig. S2**), provide important experimental controls by establishing that this system is both thermodynamically and kinetically reversible up to at least 1500 bar and that interconversion rates are solely determined by the applied pressure (and not susceptible to hysteresis effects of prior pressure applications).

As an additional verification of the suitability of pressure jumps for kinetic studies of this process, we compared the thermodynamic parameters available from an Eyring analysis of the interconversion rate. To do so, we equilibrated the protein at 1000 bar at four different temperatures between 278.1 and 291.1K, then relaxed the system to 20 bar and recorded ^13^C-edited 1D ^1^H NMR spectra to extract the apparent exchange rates from SLIP to WT (**Fig. S8**). From Eyring analysis of these data, we extracted entropic (−47.8 cal/(mol degree), or -14.2 kcal/mol at 298K) and enthalpic (7.4 kcal/mol) contributions to the activation energy, similar to values we previously obtained by monitoring re-equilibration following chromatographic isolation of the SLIP conformation ^[30]^. These data both support the interconversion via an unfolded transition state, as indicated by the large entropic barrier, and more generally indicate that the relaxation process we monitored here after pressure perturbation is the same as we previously observed with chromatographic separation ^[29, 30]^.

### Activation Volumes of the ARNT PAS-B Y456T are Comparable to the Unfolding Volumes of the WT and TRIP Variant Detected by Pressure Jump NMR

To calibrate the degree of unfolding involved in the ARNT PAS-B Y456T transition state, we compared the activation volumes of interconversion to the unfolding volumes of the WT and SLIP conformations (as adopted by the WT and TRIP variant). As neither protein substantially unfolded within our 2500 bar experimentally-accessible pressure range, we added 3 M urea to slightly lower the stability of both samples. From ^15^N/^1^H HSQC spectra we acquired at different pressures in the presence of this denaturant, it was clear that all the resolved backbone peaks showed pressure-induced reductions in intensity, with no obvious peak broadening (**Fig. S9**). This was accompanied by the appearance of intense peaks with sharp linewidths and poor ^1^H chemical shift dispersion, consistent with pressure-induced protein unfolding that completed at approximately 1750 bar (WT) and 2500 bar (TRIP) (**Fig. 4A, S9**). By fitting the protein unfolding curves for both proteins to a two-state model (**Eq. S10, Fig. 4B**), we extracted two key parameters: the average volume difference between the folded and the unfolded protein (Δ*V*_*f*_) and the free energy of folding 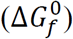 ^[20, 47]^ (**Table 1**). The unfolding volumes indicate a 43.0 ml/mol difference between the two folded structures, and the folding energies predict a 2.7:1 ratio between the SLIP and the WT conformation at ambient pressure, both similar to what we observed for ARNT PAS-B Y456T (40.5 ml/mol and 1.5:1, respectively). The activation volumes of the ARNT PAS-B Y456T, which represent the volume differences between the two folded states to the intermediate state, are smaller but comparable to the unfolding volumes of the WT and TRIP variant (−82.4 ml/mol vs. -126.3 ml/mol for WT, -37.6 ml/mol vs. -83.3 ml/mol for SLIP), reaffirming that ARNT PAS-B Y456T partially unfolds as it interconverts between conformations.

**Figure 4.**
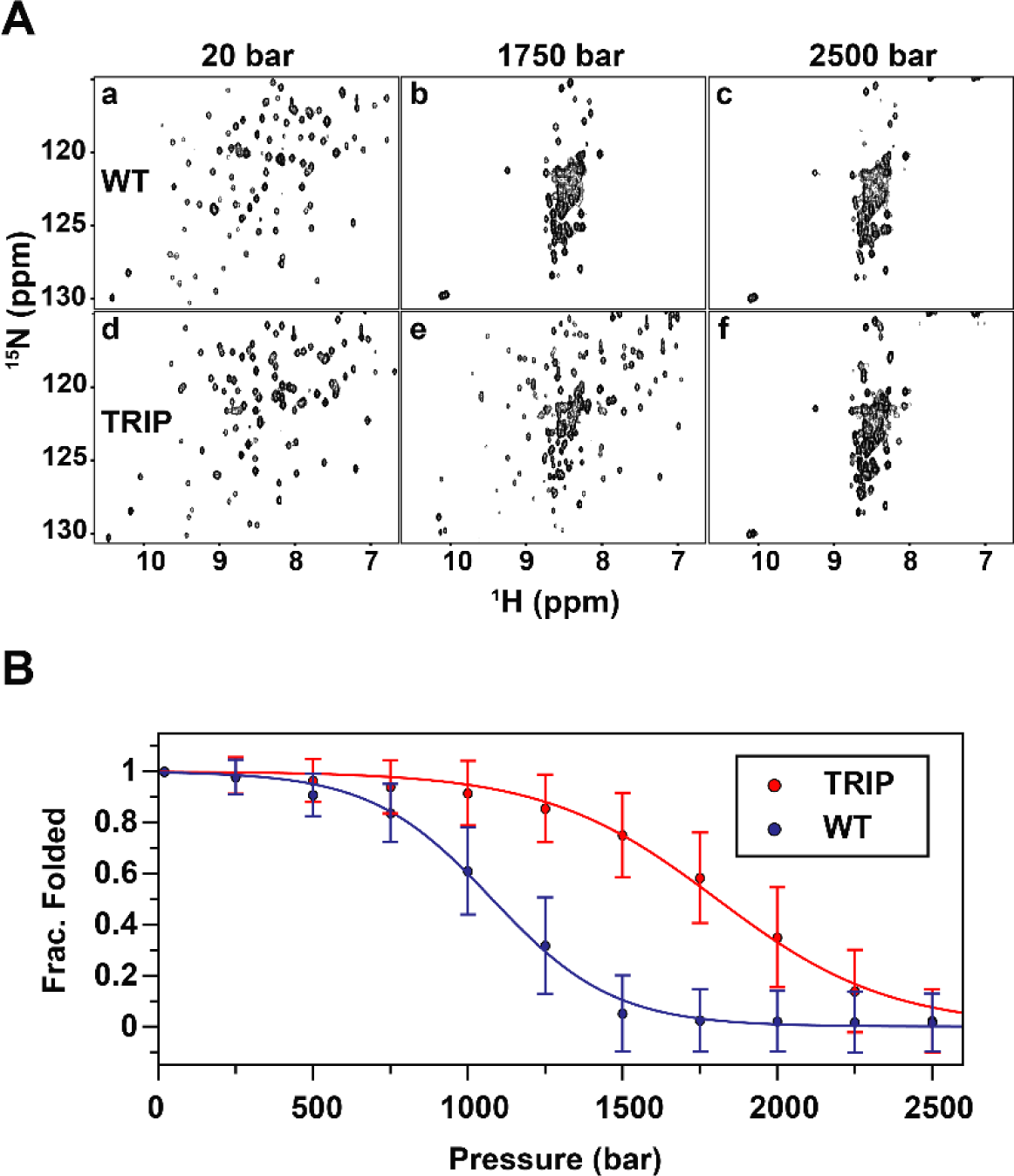
Unfolding profile of ARNT PAS-B WT and TRIP mutant detected by high-pressure NMR ^15^N/^1^H HSQC at 278 K and 3.0 M urea. A) Example ^15^N/^1^H-HSQC spectra of WT and TRIP at different pressures. (a-c) NMR spectra of WT at initial pressure (20 bar), intermediate pressure (1750 bar), and final pressure (2500 bar). (d-f) NMR spectra of TRIP mutant. B) Average unfolding curves for ARNT PAS-B WT and TRIP mutant. Peak intensities are normalized between 0 and 1.

We additionally compared the unfolding volumes of WT and TRIP to the unfolding volumes of similarly-sized proteins (10-20 kDa). Indeed, the numbers are in good agreement with several other proteins studied with pressure-dependent unfolding approaches, as summarized in **Table S1** ^[42, 45, 48-52]^. A schematic figure describing the relationships among the two folded conformations, the intermediate state, and the unfolded state is shown in **Fig. 5**.

**Figure 5.**
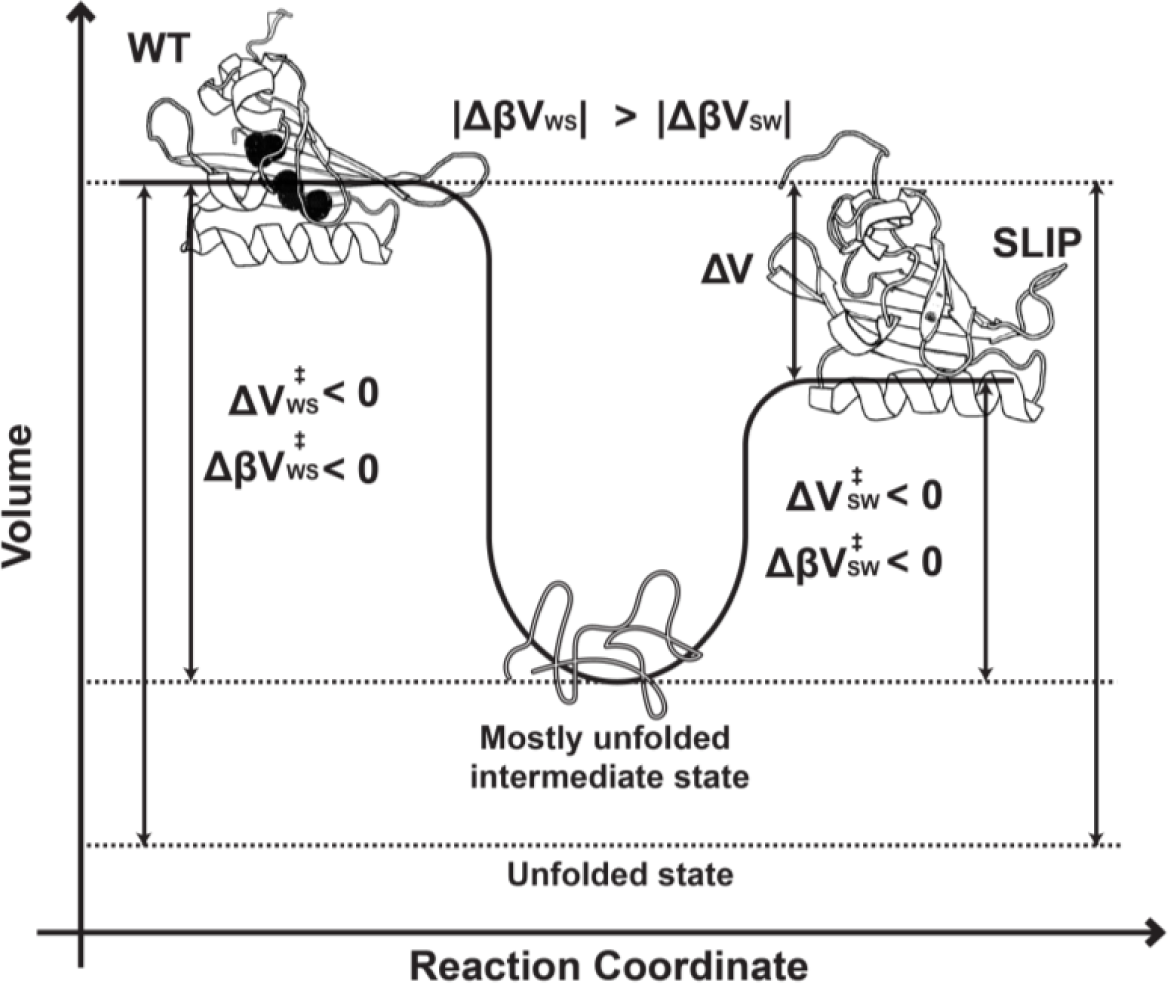
Comparison between the folded and unfolded states of ARNT PAS-B Y456T. Schematic representation comparing the folded, intermediate, and unfolded states of ARNT PAS-B Y456T. Measurement of the volumes and compressibilities are recorded in **Table 1**.

Lastly, we note that we did not include compressibility changes for the unfolding analysis of ARNT PAS-B WT and TRIP, given that these were based on global averages of signals (> 60 residues) distributed throughout both proteins which we assume to average out contributions from sites near cavities (compressibility-dependent) and those far from cavities (compressibility-independent) ^[18, 28]^.

### Pressure-induced Nonlinear Chemical Shift Changes and Residue Compressibilities Predict Cavity Locations

Certain types of pressure-induced NMR chemical shift changes have been related to conformational changes within proteins ^[26, 27]^. In particular, non-linear shift changes are hallmarks of such conformational changes, as identified by fitting chemical shifts to pressure using a linear and a nonlinear term (**Eq. S11**) ^[18]^. Of particular interest is the nonlinear coefficient (*c*_*i*_) of the backbone amide ^15^N and ^1^H nuclei, where larger values are particularly sensitive on the ability of proteins to visit multiple conformational states upon application of pressure. Such information can be analyzed in bulk, with larger average *c*_*i*_ values or broader *c*_*i*_ distributions reflecting the presence of internal cavities or voids which enable protein flexibility with increased pressure ^[18, 26, 28, 53]^. At a residue-specific level, larger *c*_*i*_ values tend to be observed from residues located near internal cavities ^[18, 28]^.

To test if this trend holds for the WT and SLIP conformations of ARNT PAS-B, we acquired ^15^N/^1^H HSQC spectra of both WT and TRIP samples of ARNT PAS-B at pressures up to 2000 bar (**Fig. 6A**). We fit the pressure-dependent changes in ^15^N chemical shifts of assignable residues to **Eq. S11**, using these to extract ^15^N *c*_*i*_ values for both conformations (**Fig. 6B**) ^[29, 31]^. While the two conformers have similar overall structures, we observed markedly different distributions of the backbone amide ^15^N *c*_*i*_ values. Specifically, we observed a global reduction of the magnitude of *c*_*i*_ in the SLIP conformation, supporting the view that it has reduced internal void volume and flexibility compared to the WT conformation. We calculated the differences in nonlinear coefficients by subtracting the absolute values of *c*_*i*_ of the SLIP conformation from the WT conformation (**Eq. S12**). Residues with the largest ^15^N *c*_*i*_ differences were located near the internal cavities or on loops of the WT structure (**Fig. 6C**), with many clustered into two regions of the protein (**Fig. 6B**). Intriguingly, the same clusters of residues have also been associated with the binding of KG-548 to ARNT PAS-B, disrupting interactions between ARNT PAS-B and the TACC3 co-activator ^[32]^. The cavities apparently collapse in the SLIP conformation due to the +3 shift and inversion of the Iβ-strand, resulting in a more packed local environment, explaining the substantial *c*_*i*_ reduction of these residues in the SLIP conformation. Interestingly, residue Y456, the site of the Y456T point mutation that creates the SLIP conformation, has one of the largest ^15^N *c*_*i*_ difference between the two conformations.

**Figure 6.**
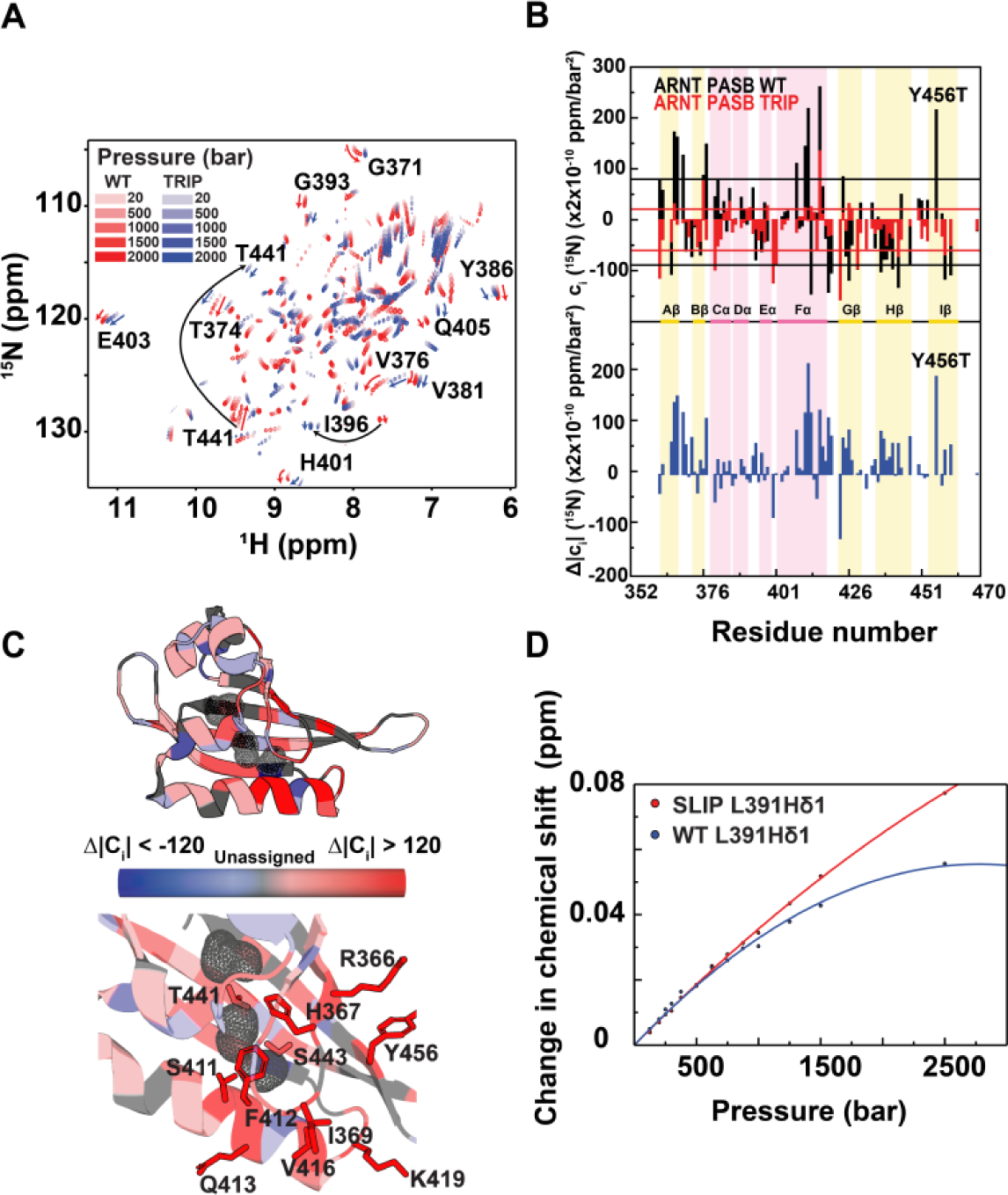
Residues surrounding cavities of ARNT PAS-B are rigidified in the SLIP conformation. A) ^15^N/^1^H HSQC spectra of WT and TRIP variant ARNT PAS-B under pressures from 20 to 2000 bar, showing large chemical shift differences between conformations despite high structural similarities between the two. Pressure-dependent non-linear chemical shift changes are also different between WT and SLIP, which are highlighted by red and blue arrows for several example residues. B) Pressure-dependent non-linear chemical shift changes of the two conformations as measured by ^15^N *c*_*i*_ (top panel) and differences in non-linearities between the two conformations (|WT *c*_*i*_|-|SLIP *c*_*i*_|) (bottom panel) mapped onto the sequence and secondary structure of WT ARNT PAS-B. The standard deviation of WT *c*_*i*_ (black lines, 84 ×2×10^−10^ ppm/bar^2^) is about 2 times the standard deviation of SLIP *c*_*i*_ (red lines, 41 ×2×10^−10^ ppm/bar^2^). C) Mapping the residue-specific differences in non-linear coefficients between WT and SLIP suggest that WT is globally more dynamic. Residues with large non-linear coefficient differences (Δ*c*_*i*_ > 120) between WT and SLIP (366, 367, 369, 411, 412, 413, 416, 419, and 456) are located near the cavities (gray mesh) or on loop regions of WT ARNT PAS-B. Other residues oriented toward cavities (i.e. T441 and S443) also show more non-linear chemical shift changes in WT than SLIP. d) ^1^H chemical shift changes of L391 Hδ1 for both the WT and SLIP conformations of ARNT PAS-B Y456T are plotted against pressure. The chemical shift change of the residue in the WT conformation shows remarkably more non-linear characteristic than it is in the SLIP conformation.

We suspected that the residue-specific compressibility also depends on whether the residue is near a cavity. As expected, when we fit the pressure dependence of the L391 Hδ1 peak to **Eq. S11**, we obtained a larger nonlinear coefficient for the WT conformation than the SLIP conformation (**Fig. 6D**), matching the compressibility difference observed between the two states, confirming our speculation. We also compared the pressure-dependent response of several other methyl peaks corresponding to the WT and SLIP conformation using the ^13^C/^1^H-HSQC spectra of ARNT PAS-B Y456T (**Fig. S10**). We found several cavity-oriented methyl groups to have markedly larger ^1^H or ^13^C *c*_*i*_ values for the WT conformation (L391 δ2, I396 δ1, L408 δ1 and δ2, and M439 ε1). Correspondingly, the pressure-dependent WT to SLIP transitions were also slowed down at higher pressures as the result of negative compressibility changes, similar to L391 δ1.

Since both nonlinear chemical shift changes and compressibility probe for the volume-dependent local environment a residue resides in, measuring the nonlinear coefficient of chemical shift changes and compressibility can complement each other for more accurate characterization of cavities in proteins (as has been linked mathematically in the past ^[54]^). This is in line with the prior notion that larger compressibility is correlated with larger volume fluctuations, which are generally observed near cavities, where residues are less tightly packed ^[55]^. Remarkably, the cavities in the WT ARNT PAS-B were not initially observed in the solution structure ^[31]^, showing the potential of pressure-NMR as the means of rapidly identifying smaller cavities from moderate resolution structures where cavities and/or ligand binding pockets are not obvious ^[31]^.

## CONCLUSION

The shift of the β-strand register at an interaction interface enabled by a single point mutation is mechanistically intriguing and raises fundamental questions about the relative stabilities of the ground state and energetically close alternate conformers ^[30, 56, 57]^. Using 1D and 2D pressure-jump NMR experiments, we examined one such case by elucidating the volume and compressibility differences between two stably folded conformations of ARNT PAS-B which interconvert when enabled by a single point mutation, Y456T. We demonstrated that the wildtype (WT) state has a substantially larger internal volume and compressibility compared to the alternate slip conformation (SLIP), and that these differences can be largely attributed to the cavities unique to the WT state. Furthermore, we show that interconversion between the two states goes through a chiefly-unfolded intermediate state that is smaller in volume and compressibility than both folded conformations. Additionally, we found that the residue-specific compressibility and nonlinear chemical shift responses to pressure can be used to predict whether residues are located close to cavities.

In summary, we have shown how varying pressure can be easily applied to reversibly shift the equilibrium of a protein between two stably folded conformations, letting us quantitatively measure structural and thermodynamic parameters otherwise difficult to access ^[20]^. We believe that this approach can be applied to other proteins which undergo large scale conformational changes, particularly with recent instrumentation advances that allow millisecond-timescale pressure jumps ^[23, 24]^. We anticipate that such studies will be particularly useful studying systems where protein flexibility is essential, including enzymatic conformational changes, protein/ligand interactions, and metamorphic systems ^[9-11]^.

## ACKNOWLEDGEMENTS

This work was supported by NSF grant MCB 1818148 (K.H.G.) and Fonds de Recherche Québec – Nature et Technologie fellowship B3X (D.G.). We thank Bruce Johnson (CUNY ASRC) for developing new extensions of the NvFX software package ^[58, 59]^ for pressure NMR analysis, and members of the Gardner laboratory for constructive comments.

