## Supporting Information for "High-Pressure NMR Reveals Volume and Compressibility Differences Between Two Stably Folded Protein Conformations"

### MATERIALS AND METHODS

#### Protein Purification and Expression

$His_6$-tagged human ARNT PAS-B (residues 356-470) WT and mutants (Y456T and F444Q/F446A/Y456T) were uniformly labeled with $^{13}C$ and $^{15}N$ and overexpressed in BL21 (DE3) E. coli. Isotopically-labeled proteins were obtained by growing cells in M9 media supplemented with 3 g/L U-$^{13}C_6$ glucose and 1 g/L $^{15}NH_4Cl$ as the carbon and nitrogen sources, respectively. Cells were incubated at 37 °C until an $OD_{600}$ of 0.6-0.8. Protein expression was induced with 0.5 mM Isopropyl-β-D-thiogalactopyranoside (IPTG) and incubated overnight (~ 16-18 hr) at 18 °C. Cells were harvested next day, cell pellets were resuspended in 50 mM Tris (pH 7.5), 150 mM NaCl, 10 mM imidazole, 1 mM DTT buffer containing 1 mM PMSF, lysed with sonication, and centrifuged. Supernatant was incubated with 5 mL of $Ni^{2+}$ Sepharose™ High Performance beads and eluted with 500 mM imidazole buffer. $His_6$-Tagged ARNT PAS-B proteins were exchanged into a low imidazole buffer (8 mM imidazole, 50 mM Tris (pH 7.5), 150 mM NaCl, and 1 mM DTT) and incubated overnight with $His_6$-tobacco etch virus (TEV) protease for tag cleavage. The freed $His_6$-tag and $His_6$-TEV were removed by a second round of $Ni^{2+}$ column purification. ARNT PAS-B proteins were further purified by passing through a Superdex 75 size exclusion column and exchanged into baroresistant buffer (44.7 mM Tris pH 7.0, 5.3 mM phosphate, 17 mM NaCl, 1 mM DTT) [1] and concentrated with an Amicon stirred ultrafiltration unit with a 3 kDa filter to 320 μM. Samples were flash frozen with liquid nitrogen and were stored at -80 °C for later use.

#### Pressure-Jump NMR Experiments

Solution NMR experiments were performed on a Bruker Avance III HD 700 MHz NMR spectrometer equipped with a 5-mm QCI-F cryoprobe. NMR samples were prepared by mixing 80% of the purified protein samples with 20% $D_2O$ yielding a final concentration of around 250 μM. Samples were loaded into a commercially-available pressure resistant NMR cell (Daedalus Innovations, Aston, PA, USA), with pressure between 20 bar to 2500 bar applied through a Xtreme-60 Syringe Pump from the same vendor. The baseline pressure was set to 20 bar as it is a low pressure that can be stably maintained by the pump. We performed two types of pressure-jump experiments, direct pressure-jump experiments where pressure was increased directly from 20 bar to the destination pressure of up to 2500 bar, and cumulative step pressure-jump experiments with constant intervals, where pressure was increased incrementally from baseline pressure to the final pressure of 2500 bar. To set up direct pressure-jump experiments, samples were thawed and stored on ice, and first equilibrated at 20 bar and 278.1 K for 1 hr and raised to a higher pressure (between 125 bar to 1500 bar) and equilibrated at that pressure. As identified in preliminary experiments, the duration of equilibration process varies depending on the pressure: interconversion of the two populations at 20 bar takes around 12 hours to complete, while it occurs more quickly as pressure increases [2]. Equilibration time under pressure was therefore set to 12 hr at 125 bar, and gradually reduced to 5 hr for pressures between 500 to 1000 bar, and finally 4 hr for pressures beyond 1000 bar. During the equilibration, a series of 1D $^{13}$C edited $^{1}$H NMR experiments were recorded to detect the rate of interconversion, with each spectrum requiring ca. 4.25 min to acquire. The system was relaxed for 12.5 hr at 20 bar post equilibrations at different pressures, 1D $^{13}$C-edited $^{1}$H NMR experiments were also recorded during relaxation. For step pressure-jump experiments, samples were first equilibrated at 20 bar and 278.1K, then pressurized and equilibrated in steps of 250 bar, until reaching 2500 bar. Equilibration time was set to 9 hr at 250 bar, 6 hr at 500 bar, 5 hr for 750 and 1000 bar, and 4 hr for 1250, 1500, 1750, 2000, and 2250 bar. Equilibration time was extended to 6 hr at 2500 bar to monitor any unfolding effects at the highest pressure safely accessible with our instrumentation. 1D NMR spectra were acquired as described earlier. $^{15}$N/$^{1}$H and $^{13}$C/$^{1}$H HSQC spectra were also acquired at system equilibrium at 20, 250, 500, 750, 1000, 1250, 1500, and 2500 bar. $^{15}$N/$^{1}$H HSQC experiments were acquired with 16 scans and FIDs of 2048x256 complex points. $^{13}$C/$^{1}$H HSQC experiments were acquired with 8 scans and FID of 2048x256 complex points. All NMR spectra were processed and analyzed with NMRFx Analyst and NMRViewJ [3, 4]. Chemical shift assignments from previous NMR assignments of ARNT PAS-B WT and F444Q/F446A/Y456T mutant were used for all analyses [2, 5].

Unfolding measurements of ARNT PAS-B WT and the F444Q/F446A/Y456T variant under pressures accessible by our apparatus (20-2500 bar) was achieved by the addition of urea (up to 3.0 M). $^{15}$N/$^{1}$H HSQC spectra were acquired between 20 and 2500 bar with increments of 250 bar. Peak assignments from previous work were obtained [5, 6], and transferred to the acquired spectra following urea titration experiments between 0.5 and 3.0 M.

**NMR Data Analysis**

In the 1D $^{13}$C-edited $^{1}$H NMR experiments, two peaks corresponding to the L391-δ1 methyl group of the WT and SLIP conformations were used to determine the populations of each conformational state of the protein in the sample. The relative populations (SLIP/WT) at different pressures were obtained by taking the ratio of the integrals of the two peaks. The equilibration (change in relative population) under different pressures and the rates of interconversion were obtained by plotting the relative population as a function of time.

The equilibrium constant K between the two conformers of the ARNT PAS-B Y456T is given by the relation

$$K = \frac{[SLIP]}{[WT]} = e^{-\Delta G/RT} \qquad \text{(S1)}$$

Since the temperature T was kept constant during the pressure jump experiments, the free energy $\Delta G$ scales only with pressure p. The free energy as a function of pressure can be approximated by a Taylor expansion

$$\Delta G(p) = \Delta G^0 + \Delta V^0 (p - p_0) - \frac{1}{2} \times \Delta\beta V^0 (p - p_0)^2 + \ldots \frac{\Delta G^n(p_0)}{n!}(p - p_0)^n \qquad \text{(S2)}$$

$\Delta G^0$ is the free energy between the two conformers at ambient pressure, $\Delta V^0$ is the volume difference between the two conformers, representing the first derivative of free energy with respect to pressure, and compressibility between the two conformations, $\Delta\beta V^0$, is the second derivative of free energy, with respect to pressure. The equilibrium constants were plotted as a function of pressure, and fit to the following equation, incorporating up to the third term of the Taylor series [7]:

$$K_{eq} = e^{-\left(\Delta G^0 + (p - p_0)\Delta V^0 - 1/2 \times \Delta\beta V^0 (p - p_0)^2\right)/RT} \qquad \text{(S3)}$$

The pressure was referenced to atmospheric pressure, $p_0$=1 bar.

The observed rate of increase of one conformation (or decrease of the other conformation) as a function of pressure could be fit to a biexponential equation:

$$k_{obs} = k_{sw0} e^{-\left(p\Delta V_{sw}^{\ddagger} - 1/2 \Delta\beta V_{sw}^{\ddagger} p^2\right)/RT} + k_{ws0} e^{-\left(p\Delta V_{ws}^{\ddagger} - 1/2 \Delta\beta V_{ws}^{\ddagger} p^2\right)/RT} \qquad \text{(S4)}$$

yielding the activation volumes ($\Delta V_{sw}^{\ddagger}$ and $\Delta V_{ws}^{\ddagger}$) and compressibility differences between the two folded conformations (WT and SLIP) to the previously-proposed, chiefly unfolded intermediate state ($\Delta\beta V_{sw}^{\ddagger}$ and $\Delta\beta V_{ws}^{\ddagger}$) [2]. Together with the two initial rate constants at ambient pressure

($k_{sw0}$, SLIP to WT, and $k_{ws0}$, WT to SLIP), there were 6 unknown parameters to be fit. To better approximate the two exponential terms, rates at different pressures were solved numerically using the following differential equation pair:

$$\frac{d[S]}{dt} = -k_{sw}[S] + k_{ws}[W] \qquad \text{(S5)}$$

$$\frac{d[S]}{dt} = k_{sw}[S] - k_{ws}[W] \qquad \text{(S6)}$$

The fittings were done in R, using the package deSolve as the ordinary differential equation solver, and Levenberg-Marquardt Nonlinear Least-Squares Algorithm in minpack.lm for parameter fitting [8].The uncertainty of the fitting was estimated by calculating the 95% confidence intervals from the variance covariance matrix of the fit rate constants. 95% confidence intervals were further verified with a bootstrapping procedure. Random noises with mean of 0 and variance of the mean square error from the model were added to the raw data and fit repeatedly. The estimated parameters were computed to determine if they fall within the 95% confidence interval. $k_{sw}$ and $k_{ws}$ obtained were plotted separately as functions of pressure, and fit individually to:

$$k_{sw} = k_{sw0} e^{-\left(p\Delta V_{sw}^{\ddagger} - 1/2\Delta\beta V_{sw}^{\ddagger} p^2\right)/RT} \qquad \text{(S7)}$$

$$k_{ws} = k_{ws0} e^{-\left(p\Delta V_{ws}^{\ddagger} - 1/2\Delta\beta V_{ws}^{\ddagger} p^2\right)/RT} \qquad \text{(S8)}$$

obtaining rates at ambient pressure, activation volumes, and activation compressibilities as described earlier.

The temperature dependence of the relaxation rates was calculated as the apparent rates during the 1000 to 20 bar re-equilibration step at temperatures of T = 278.1, 283.1, 288.1, and 291.1 K. The change of relative population ([SLIP]/[WT]) over time was fit to a single exponential with the apparent rate of

$$k_{app} = k_{ws} + k_{sw} \qquad \text{(S9)}$$

An Eyring relationship between rate and temperature was found by plotting ln($k_{app}$/T) against 1/T. The plot was fit to a linear equation to extract the entropic and enthalpic contributions of activation energy of transition from SLIP to WT during relaxation at 20 bar.

For the pressure-dependent unfolding experiments, $^{15}$N/$^{1}$H HSQC spectra at different pressures were collected, maximum peak intensities in each spectrum were obtained and compared with the peaks corresponding to the same residues collected at different pressures. Overlapping peaks were excluded from the analysis. The intensity averages at different pressures were plotted as a function of pressure, and fit to a two-state unfolding model with XMGrace [9],

$$\frac{I}{I_0} = \frac{e^{-\left(\Delta G_f^0 + P\Delta V_f\right)/RT}}{1 + e^{-\left(\Delta G_f^0 + P\Delta V_f\right)/RT}} \qquad \text{(S10)}$$

where $I_0$ is the peak intensity at initial (20 bar) pressure, R is gas constant, and T is temperature. The apparent volume difference between the folded and unfolded state ($\Delta G_f^0$) and the free energy of folding ($\Delta V_f$) were extracted. For simplicity, the max and min plateau values were set to 1 and 0.

For the pressure-jump $^{15}N/^{1}H$ HSQC experiments, we analyzed the peaks under different pressures and determined the change in chemical shifts and intensity under pressures compared to the reference chemical shift (positions of peaks at 20 bar). We fit the change in chemical shifts to the following equation to determine the nonlinear coefficients[7, 10, 11]:

$$\Delta\delta_i = b_i\Delta p + c_i\Delta p^2 \qquad \text{(S11)}$$

Where p is pressure (bars), $b_i$ (parts per million per bar) and $c_i$ (parts per million per square bars) are the first and second order pressure dependence coefficients on chemical shifts for the ith residue. A large $c_i$ value suggests high nonlinear response to pressure. Fittings were performed with the built-in pressure-analysis function of NMRViewJ [3]. Residue-specific nonlinear coefficient differences between the two conformations were calculated by subtracting the absolute values of $c_i$ corresponding to each conformation:

$$\Delta c_i = \left|c_{iwt}\right| - \left|c_{islip}\right| \qquad \text{(S12)}$$

Values were mapped to the crystal structure of WT ARNT PAS-B [12] and visualized with PyMOL (The PyMOL Molecular Graphics System, Version 2.0 Schrödinger, LLC).

**Void Volume Calculation**

Void volumes of ARNT PAS-B WT and SLIP conformations were calculated with the ProteinVolume software package by Chen and Makhatadze [13]. Briefly, the total solvent-excluded volume of the protein is calculated by filling the space within the protein's molecular surface with 0.02 Å probes. The van der Waals volume is calculated with the same process but only counting the probes that are within the van der Waals radius of any atoms. The void volume $V_{void}$ is given by the difference between the two volumes calculated. Solution NMR structures of

WT and SLIP were used for the calculation [5, 6], using the average void volume of all 20 conformers in these ensembles.

#### SUPPLEMENTARY TABLES

#### Table S1. Unfolding Volumes of Proteins with Similar Size as ARNT PAS-B

| Protein | Size (kDa) | Unfolding Volume (ml/mol) | Temperature (K) | Method | Reference |
|---|---|---|---|---|---|
| **Staphylococcal nuclease** | 17 | −90 ± 15 | 275 | High-Pressure Fluorescence Measurements | [14] |
| | | −87 ± 20 | 278 | FTIR | |
| **Δ + PHS (SNase Stable vatiants)** | 17 | −80 ± 8 | 293 | NMR | [15] |
| **I92A** | | −148 ± 21 | | - | |
| **L125A** | | −108 ± 3 | | - | |
| **V66A** | | −104 ± 10 | | - | |
| **L103A** | | −106 ± 4 | | - | |
| **Ribonuclease A** | 14 | −46.5 ± 3.3 | 313 | UV-Vis | [16] |
| **CTL9 WT** | 10 | −82.95 | 288 | NMR | [17] |
| **CTL9 I98A** | - | -108.91 | | - | |
| **Nank 1-7** | ~26 | −44 ± 4.1 | 293 | High-pressure fluorescence measurement | [18] |
| **Nank 1-6** | | −48 ± 5.3 | | - | |
| **Nank 1-5** | | −47 ± 9 | | - | |
| **Nank 2-7** | | −55 ± 4.7 | | - | |
| **Nank 3-7** | | −51 ± 5.58 | | - | |
| **Nank 4-7** | | −39.4 ± 5 | | - | |
| **Metmyoglobin pH 5.15** | 17 | -117 | 278 | High-pressure optical bomb | [19] |
| **Metmyoglobin pH 5.75** | - | -127 | | - | |

#### SUPPLEMENTARY FIGURES

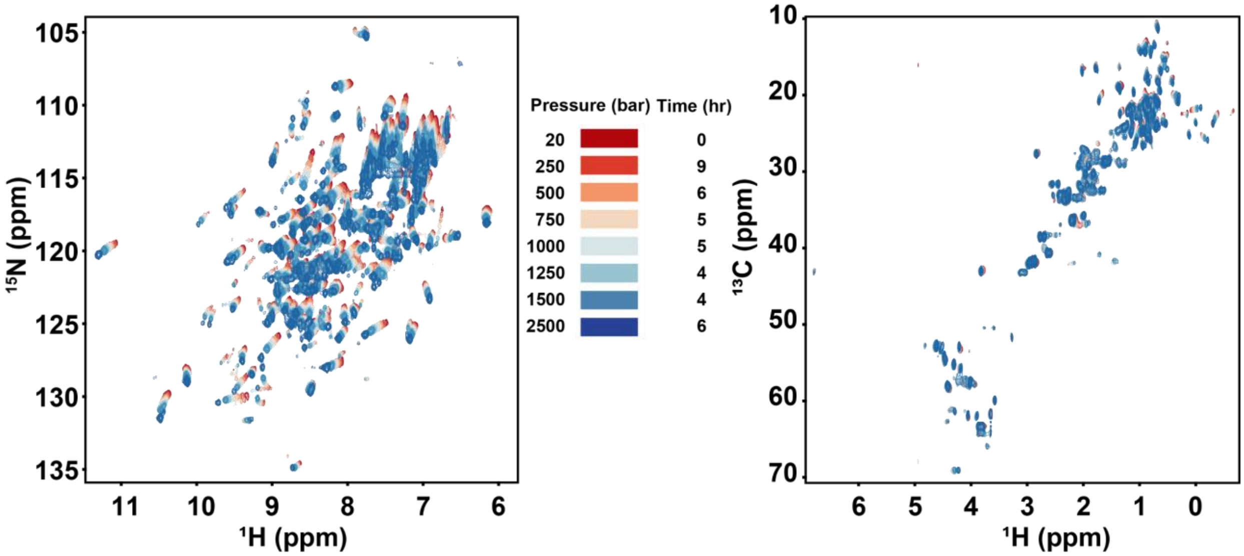

**Fig. S1. Overlaid 15N/1H and 13C/1H HSQC spectra of ARNT PAS-B Y456T recorded over different pressures.** Spectra were recorded after the system was equilibrated under 20, 250, 500, 750, 1000, 1250, 1500, 2500 bar pressure at 278.1K. Most peaks in $^{15}$N/$^{1}$H HSQC spectra (left panel) show discernable pressure-dependent chemical shift and intensity changes. Peaks in $^{13}$C/$^{1}$H HSQC spectra (right panel) also show pressure-dependent changes in chemical shift and intensity, but to a lesser extent. Most discernable changes in $^{13}$C/$^{1}$H HSQC spectra are from methyl signals. Times indicated are the time the system was equilibrated at each pressure before acquiring the HSQC spectrum.

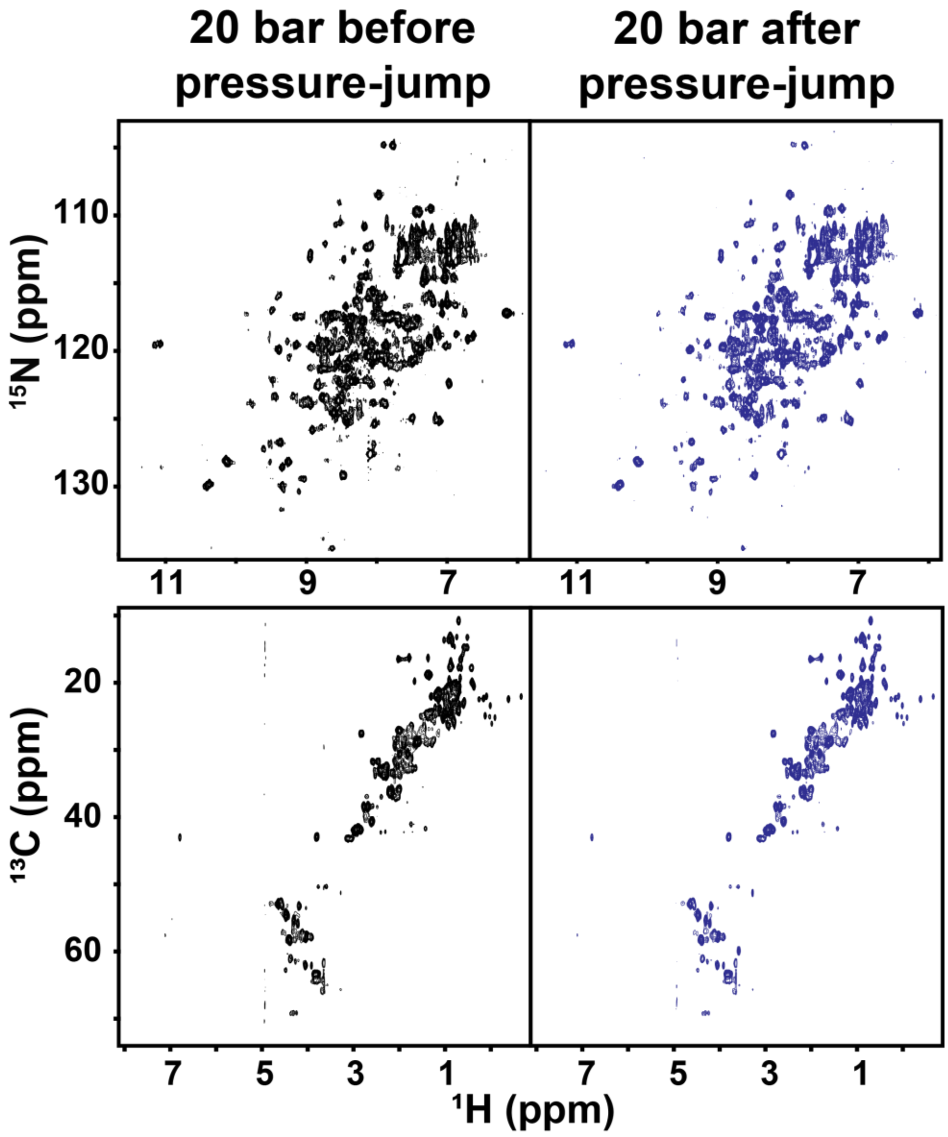

**Fig. S2. Comparison of 15N/1H and 13C/1H HSQC spectra recorded at 20 bar pre- and post- pressure-jump NMR experiments.** $^{15}N/^{1}H$ (top) and $^{13}C/^{1}H$ (down) HSQC spectra of ARNT PAS-B Y456T were collected at 20 bar pressure before (left, black) and after (right, blue) going through culminative pressure-jump experiments (sample was pressurized from 250 bar to up to 2500 bar, before re-equilibrated at 20 bar). The spectra show no significant differences, implying pressure-induced chemical shift and intensity changes are reversible for at least up to 2500 bar.

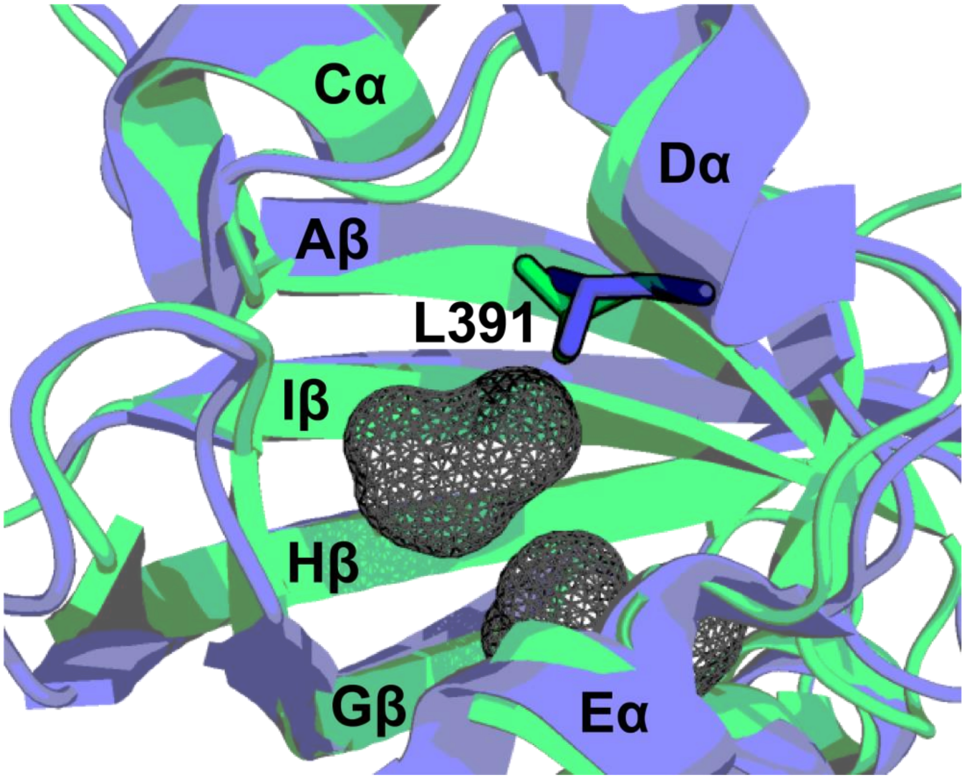

**Fig. S3. Structural differences between the WT and SLIP conformation of ARNT.** Superimposed WT (blue, PDB ID: 4EQ1[20]) and SLIP (green, PDB ID: 2K7S [5]) structures with major structural features labeled. The view is zoomed in at the internal cavities of the WT conformation, with the side chain of L391 highlighted.

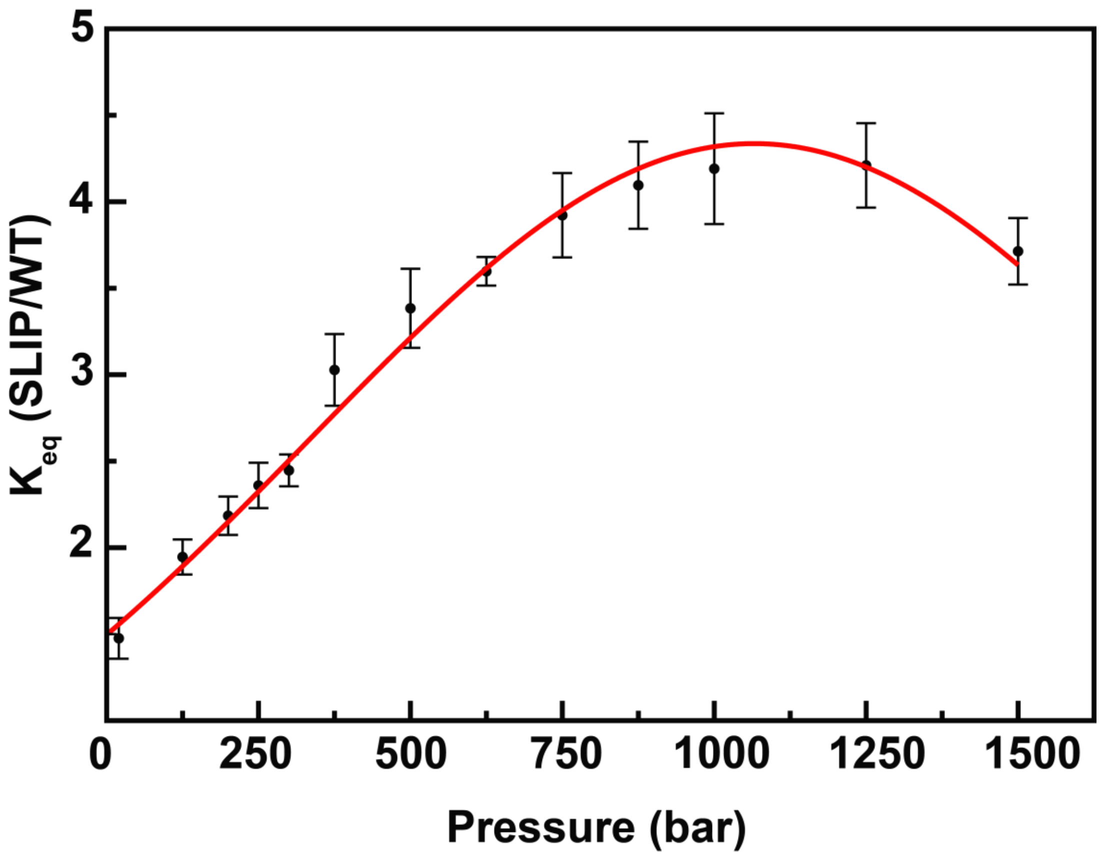

**Fig. S4. Equilibrium constant of ARNT PAS-B Y456T (SLIP/WT) fitted as a function of pressure.** Pressure of the system was individually jumped from 20 bar to various pressures (up to 1500 bar). Interconversion of the two L391 H$\delta$1 peaks corresponding to the two conformations were monitored to obtain the equilibrium constants at different pressures. Quantification of volume difference between the two conformations ($\Delta V$, SLIP-WT) and compressibility difference ($\Delta\beta V$, SLIP-WT) were obtained from the fitting.

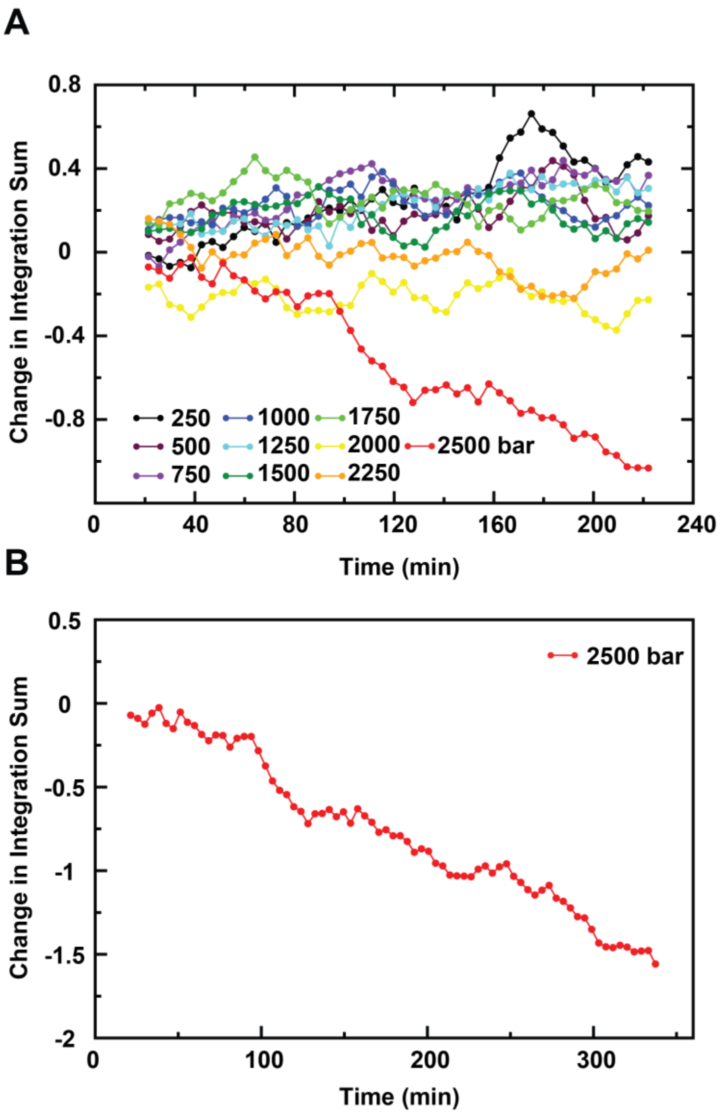

**Fig. S5. Fluctuation of the total population of the WT and SLIP conformations at different pressures.** A**)** The change in integral sum (referenced to the average of the first three data points) of the two peaks corresponding to the WT and SLIP conformations were monitored during system equilibration at different pressures. 7-point averages are plotted against time. B) 7-point averages of integral sum fluctuation at 2500 bar is plotted for extended time (6 hours).

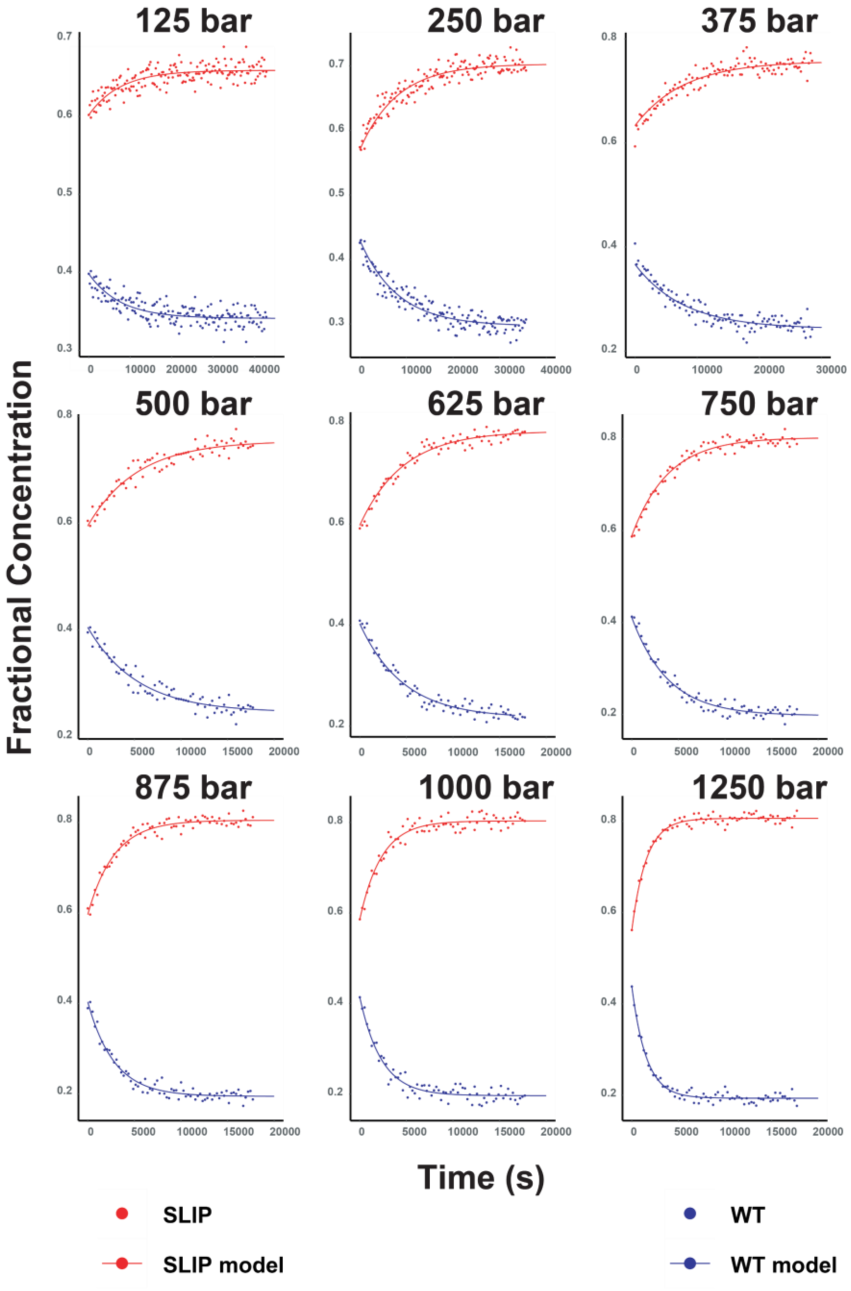

**Fig. S6. Numerical solutions to rate equations under pressure.** Rate constants $k_{sw}$ and $k_{ws}$ are estimated by solving the ordinary differential equation pairs (see Material & Methods for details). Plotted are the fractional concentration of WT (blue) and SLIP (red) over time. Solid lines are the model with the estimated rate constants.

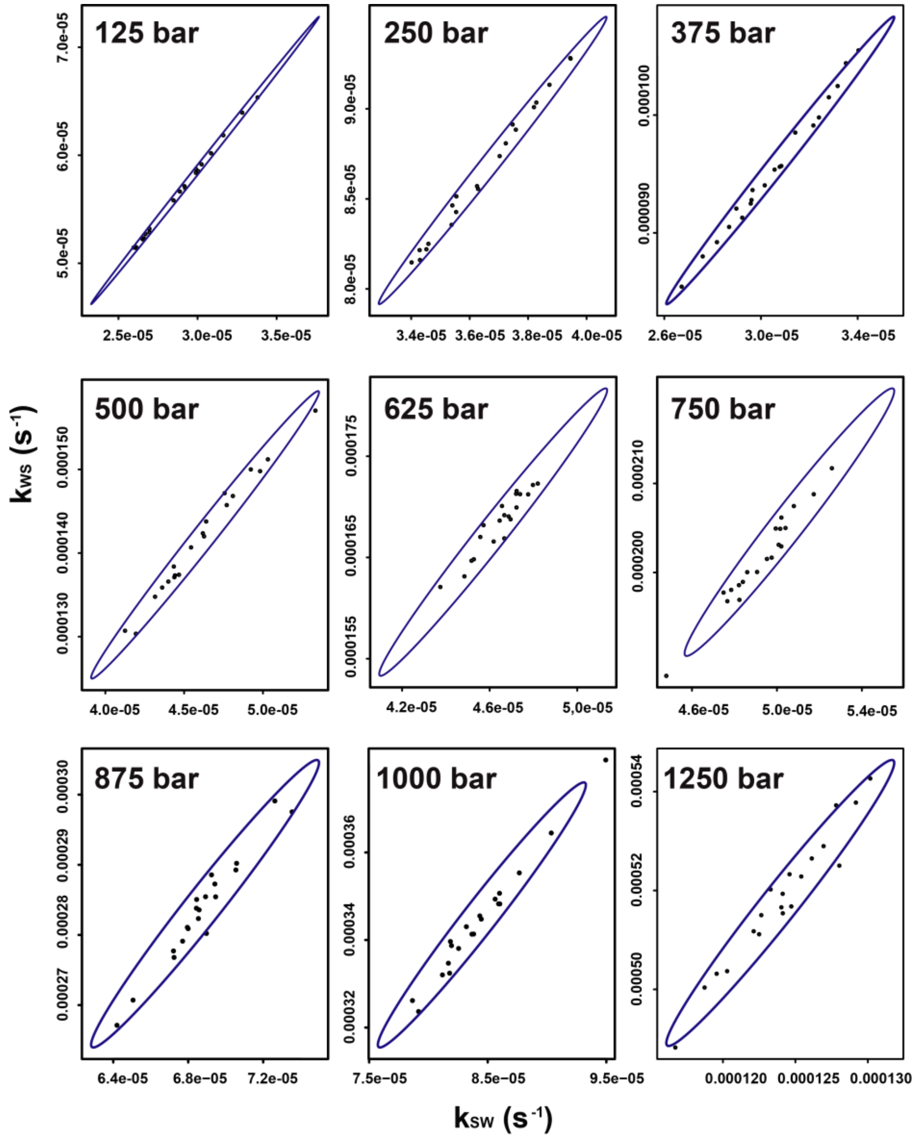

**Fig. S7. Estimation of the parameter uncertainties.** 95% confidence ellipsoids are calculated and plotted based on the variance covariance matrix of the estimated rates ($k_{ws}$ and $k_{sw}$). To validate confidence interval, uncertainties were estimated again using a bootstrapping process. 20 datasets were generated by adding random normal noise to the original data points and fitted again. The newly fitted parameters were plotted onto the ellipsoid plots. Most of these fits (~ 95%) indeed fall within the confidence interval). The plots also show strong correlations between the two rates, the pressure-dependent equilibrium constants at different pressures may be estimated from the slops of the plots.

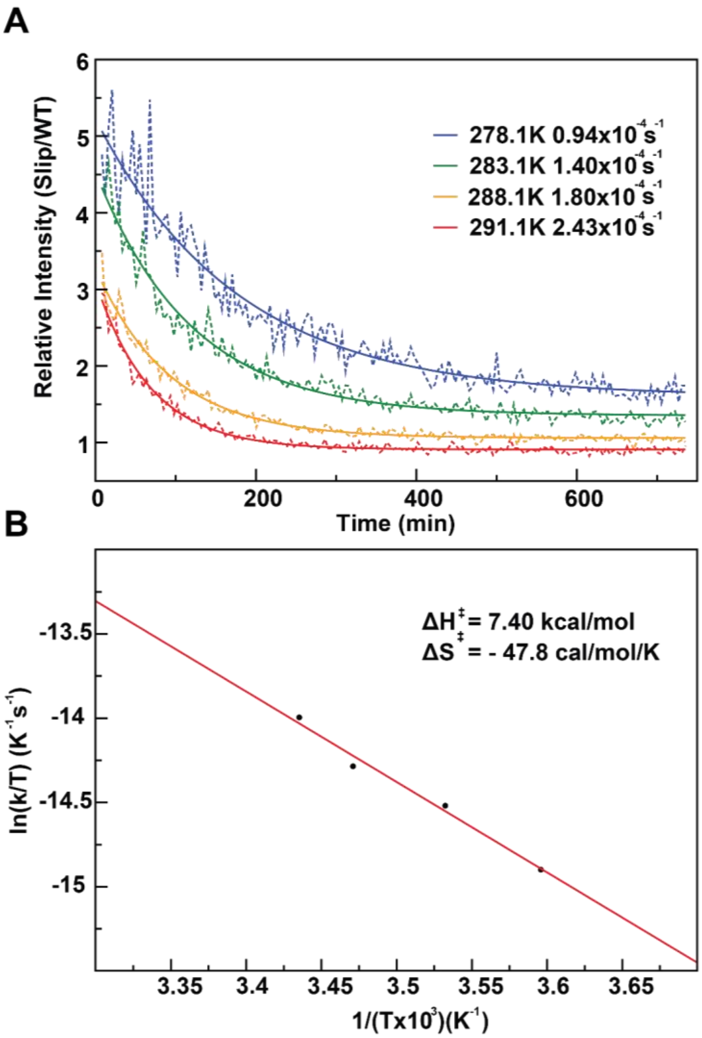

**Fig. S8. Temperature dependence of ARNT PAS-B Y456T interconversion between the SLIP and WT conformations.** A) ARNT PAS-B Y456T was equilibrated at 1000 bar and relaxed at 20 bar. $^{13}C$ edited 1D NMR spectra were collected during relaxation to monitor relative population (SLIP/WT) change over time. Exchange rates are extracted from single exponential fittings at 4 different temperatures (278.1K, blue; 283.1K, green; 288.1K, yellow; 291.1K, red). B) Interconversion between the WT and SLIP conformations at 20 bar follows a linear Eyring dependence. Activation enthalpy and entropy are extracted from the Eyring plot.

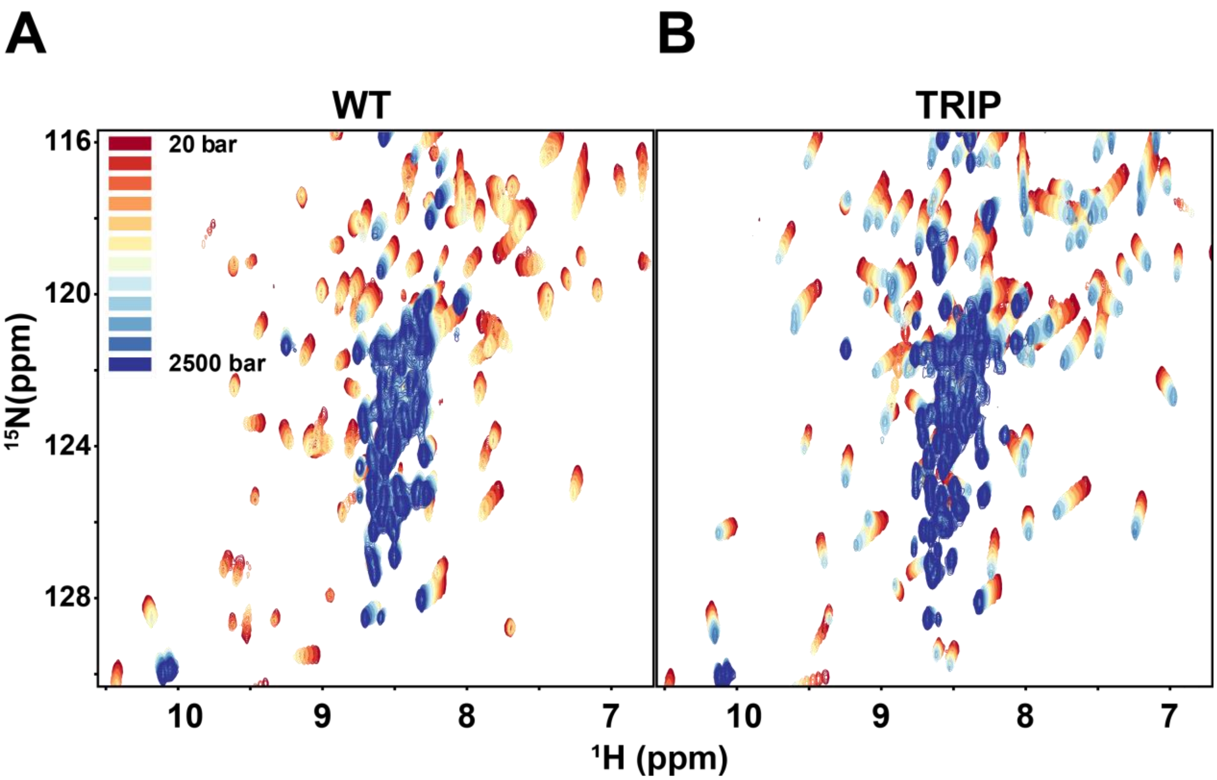

**Fig. S9. 15N/1H HSQC spectra of ARNT PAS-B WT and TRIP variant under pressure at 278 K and 3.0 M urea.** A) Superimposed WT protein spectra from 20 bar to 2500 bar with 250 bar steps (20, 250, 500, 750, …, up to 2500 bar). B) Superimposed TRIP mutant from 20 bar to 2500 bar with 250 bar steps.

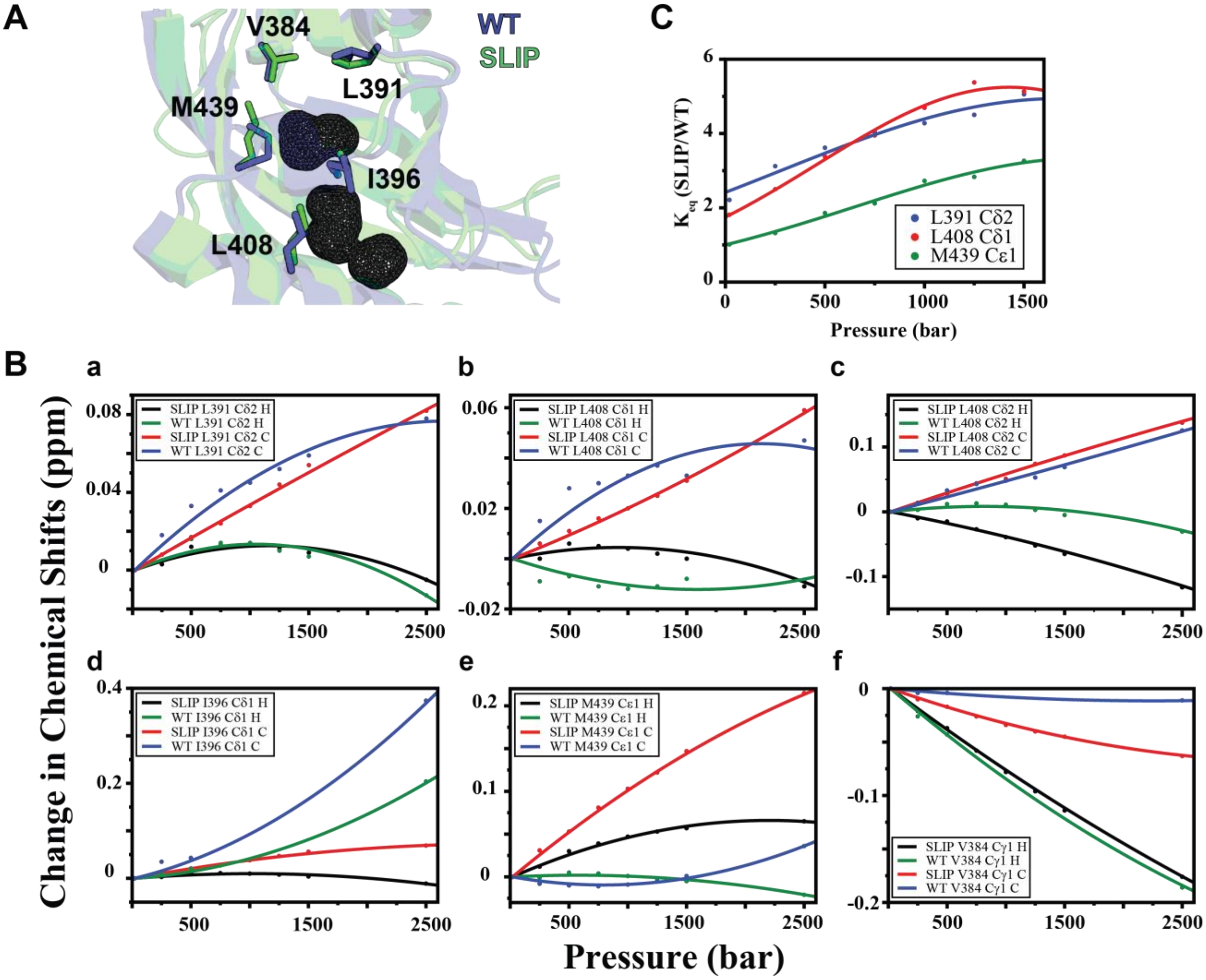

**Fig. S10. Pressure-dependent chemical shift changes and equilibrium constants (SLIP/WT) of cavity-oriented residues of ARNT PAS-B Y456T.** A) Residues representing the WT (slate blue) and SLIP (lime green) conformations suitable for analysis are highlighted and labeled. Cavities uniquely exist in the WT state are shown as gray wireframe surfaces. B) Chemical shift changes of side chain methyl groups show great variations between the WT and the SLIP conformations, with more non-linear characteristics in the WT conformation. Curves representing I396 Cδ1 and V384 Cγ1 are fitted with less data points as signals at some pressures are overlapped with other peaks, rendering them unassignable. C) Equilibrium constant of three side chain methyl groups (probed by $^{13}C/^{1}H$-HSQC) are plotted against pressure. All three cavity-oriented methyl groups show similar pressure-dependent behavior as L391 Hδ1.
